# Single-Cell RNA Sequencing Reveals Distinct Cellular and Molecular Features of Steroid-Induced Osteonecrosis of the Femoral Head Compared to Alcohol-Induced Osteonecrosis and Hip Osteoarthritis

**DOI:** 10.64898/2026.11.01.763474

**Authors:** Arijita Sarkar, Matthew C. Gallo, Jennifer A. Bell, Cory K. Mayfield, Jacob R. Ball, Fergui Hernandez, Elizabeth Lechtholz-Zey, Tara Shelby, Osamu Sugiyama, Denis Evseenko, Jay R. Lieberman

## Abstract

Osteonecrosis of the femoral head (ONFH) remains a devastating complication of glucocorticoid therapy and alcohol abuse, yet the pathophysiology is poorly understood. Using single-cell RNA sequencing, we analyzed femoral head specimens from patients undergoing total hip arthroplasty due to steroid-induced ONFH, alcohol-induced ONFH, or hip osteoarthritis (OA). Our comprehensive analysis across conditions revealed etiology-specific cellular compositions and transcriptional profiles. Steroid-induced ONFH demonstrated a distinctive immunological signature characterized by a neutrophil-predominant, elevated innate-to-adaptive immune cell ratio compared to alcohol-induced ONFH or OA. T cell analysis revealed impaired maturation in steroid-induced ONFH, with abundant naïve T cells but reduced mature CD4+ T cells and CD4+ memory T cells compared to both alcohol-induced ONFH and OA. In addition, compared to alcohol-induced ONFH, steroid-induced ONFH was associated with an increase in exhausted/dysfunctional CD8+ T cells which suggests an impaired response to stress and reflects chronic immune dysfunction. Within stromal cell populations, steroid-induced ONFH demonstrated an increase in pericytes, endothelial cells, and mesenchymal cell with osteogenic features as compared to both alcohol-induced ONFH or OA. These findings collectively reveal novel cellular mechanisms underlying steroid-induced ONFH which may contribute to further investigation of etiology-specific pathogenesis and treatment strategies.

**LAY SUMMARY:** Osteonecrosis of the femoral head is a painful condition where bone tissue in the hip joint dies, often associated with steroid medications or alcohol use. We analyzed the gene expression of thousands of cells from hip tissue samples to understand how these two risk factors contribute to bone death, and how they compare to hip degeneration seen in osteoarthritis. Each condition is characterized by unique patterns of immune response and cellular dysfunction. Despite causing similar hip problems, steroids and alcohol may damage bone through unique biological pathways, suggesting that patients may benefit from different treatments depending on the underlying cause.

## INTRODUCTION

Osteonecrosis of the femoral head (ONFH) is a progressive disease characterized by a disruption in osseous circulation and impairment of normal cellular function, which ultimately leads to bone necrosis, femoral head collapse, and destruction of the hip joint.^1^ ONFH represents a significant healthcare challenge, with an incidence of approximately 20,000 cases per year in the United States.^1^ ONFH is the underlying diagnosis for 10% of all total hip replacements.^1^ Among the various etiologies, steroid- and alcohol-induced ONFH represent the two most common nontraumatic risk factors, accounting for up to 80% of cases.^2^ At present, the pathogenesis of ONFH remains incompletely understood.^2,3^

ONFH has been attributed to multiple pathophysiological mechanisms including endothelial cell dysfunction, direct cytotoxicity on bone-forming cells, and aberrant lipid metabolism.^4–6^ However, these discrete hypotheses have failed to capture the complex multicellular interactions that underlie disease development. Furthermore, although steroid-and alcohol-induced ONFH share similar clinical presentations and endpoints, emerging evidence suggest distinct pathogenic mechanisms may be involved.^7–9^

A critical gap in our understanding relates to the complex cellular microenvironment within the femoral head in ONFH. Traditional histopathological studies have been limited by their inability to provide comprehensive characterization of the diverse cell populations and their functional states.^8,10^ Moreover, comparative analyses between steroid-induced ONFH, alcohol-induced ONFH, and other hip pathologies such as osteoarthritis (OA) have been scarce, hindering our ability to identify etiology-specific cellular and molecular targets.^7,11^

Single-cell RNA sequencing (scRNA-seq) has emerged as a powerful technology that enables a detailed examination of cellular heterogeneity and functional states within tissues.^6,12–23^ Recently, Liao et al. utilized scRNA-seq to characterize alcohol-induced ONFH, revealing aberrant stromal cell populations and pathogenic endothelial cells compared to patients with femoral neck fractures or OA.^11^ However, no comparable analysis has been performed for steroid-induced ONFH, despite its prevalence and clinical significance.

In this study, we utilized scRNA-seq to comprehensively characterize and compare the cellular compositions and transcriptional profiles of femoral head specimens from patients with steroid-induced ONFH, alcohol-induced ONFH, and OA. Through detailed analysis of immune, vascular, and stromal cell populations, we identified multiple novel features specific to steroid-induced ONFH.

## METHODS

This study protocol was approved by the USC Health Sciences Campus & LA General Medical Center Institutional Review Board (Protocol HS-22-00552). The entire study was conducted in accordance with USC Health Sciences Campus & LA General Medical Center Institutional Review Board (Protocol HS-22-00552) guidelines. An exemption for informed consent was not provided as the tissue used for analysis is considered medical waste. The USC Health Sciences Campus & LA General Medical Center Institutional Review Board (Protocol HS-22-00552) granted a full waiver of informed consent because only de-identified discarded tissue was collected, with no patient contact, and only limited demographic information (age, sex, and relevant comorbidities) was recorded without access to identifiable patient data.

### Sample Collection Cell Isolation

Femoral head specimens were obtained from patients undergoing total hip arthroplasty for steroid-induced ONFH (n=2) or hip OA (n=3). Patient characteristics are summarized in Supplementary Table 1. Alcohol-induced ONFH femoral head sample data were obtained from publicly available data (SRA database -SRP361778) from Liao et al.’s study (n=6).^11^ To date, transcriptomic profiling has not been conducted on femoral head samples from disease-free controls. Therefore, osteoarthritis was chosen as a control condition to distinguish ONFH-specific changes from general end-stage hip degeneration that occurs in both ONFH and OA.

Following surgical resection during total hip arthroplasty, femoral head specimens were immediately processed for single-cell isolation. Each femoral head was washed once with phosphate-buffered saline (PBS). The femoral head was then sectioned along the coronal plane using an oscillating saw to expose the full depth of the femoral head. Remaining articular cartilage was removed with a rongeur. Morselized bone tissue was then systematically collected from the entire cross-section of the femoral head using a curette and rongeur, encompassing tissue from the necrotic zone, sclerotic zone, healthy margin, and subchondral bone. Approximately 20-25g (wet weight) of processed tissue was divided into 4-5 tubes (5g per tube) for enzymatic digestion.

Cells were prepared using a previously established protocol.^11^ Briefly, tissue fragments were digested in a solution containing 0.2% collagenase type II (Sigma-Aldrich) and 0.2% dispase II (Sigma-Aldrich) in serum-free Dulbecco’s Modified Eagle Medium (DMEM) at 37°C for 4 hours with continuous agitation. The digested suspension was then filtered through a 70-μm strainer, and the resulting filtrate was centrifuged at 400 × g for 5 minutes. The cell pellet was resuspended in 20 ml of serum-free DMEM.

To remove bone debris and enrich for nucleated cells, the cell suspension was subjected to density gradient centrifugation using a Percoll solution (Sigma-Aldrich). The suspension was carefully layered onto 20 ml of Percoll solution and centrifuged at 450 × g for 20 minutes with minimal acceleration and deceleration. The nucleated cell layer was collected, diluted with serum-free DMEM, and centrifuged at 400 × g for 5 minutes. RBC lysis was performed using a commercial buffer (Lonza, Basel, CH). Cells were then stained with anti-RBC antibody according to manufacturer’s protocol (BioLegend, San Diego, CA, USA), washed, and resuspended in PBS with 1% FBS containing DAPI. Fluorescence-activated cell sorting (FACS) analysis was performed to identify and sort live cells using a BD LSR II flow cytometer (BD Biosciences, Franklin Lakes, NJ, USA). The gating strategy for FACS sorting has been provided in Additional File 1.

### Single-Cell RNA Sequencing

Single-cell samples were prepared using the Chromium Next GEM Single Cell 3^/^ Reagent Kits v3 and Chip Kits (10x Genomics, Pleasanton, CA, USA), following the manufacturer’s protocol.^24^ In brief as described previously^25^, FACS-sorted live cells were resuspended in 0.04% BSA-PBS solution. Cell viability was assessed before single-cell library preparation, and only samples with viability ≥90%, were processed for scRNA-seq. Cell encapsulation and barcoding were carried out using a microfluidic droplet-based system (Chromium, 10x Genomics), generating gel bead-containing emulsions. Complementary DNA synthesis occurred within individual droplets, followed by emulsion disruption and recovery of barcoded cDNA. Purified cDNA was obtained through magnetic bead–based cleanup and subsequently amplified. Library construction involved adapter incorporation into the amplified material, additional PCR enrichment, and bead-based size selection. Library quality was evaluated using a TapeStation D1000 instrument (Agilent, Santa Clara, CA, USA), while DNA concentrations were determined using a fluorescence-based high-sensitivity assay (Qubit; Thermo Fisher Scientific, MA, USA). Final libraries were normalized, combined at equal molar ratios, and sequenced using a NextSeq system (Illumina, San Diego, CA, USA) at the institutional genomics core facility. Approximately 8000-10000 cells per sample were captured and sequenced at a depth of 20,000– 50,000 reads per cell.

### Single-Cell RNA-Sequencing Data Analysis

The raw fastq files were processed and analyzed as described previously ^25^. In brief, raw fastq files were checked for their quality, trimmed, and reads with an average base quality score per position > 30 was considered for further analysis using Cell Ranger (v7, 10x Genomics). The raw sequencing data were demultiplexed to assign reads to individual cells based on their unique cell barcodes and unique molecular identifiers (UMIs). The reads were then aligned using STAR aligner (v2.7.3)^26^ to the reference human genome (GRCh38), and a gene expression matrix was generated, quantifying the number of transcripts detected for each gene in each cell. Quality control measures were implemented using the DropletUtils package (v1.16.0) to assess the quality of the cells.^27,28^ Cells expressing fewer than 200 genes or more than 6,000 genes, as well as cells with >10% mitochondrial gene expression, were removed during quality control were excluded from further analysis. To identify and remove doublets, Doubletfinder (v2.0.3) was employed.^29^ Cells predicted to be doublets were filtered out to enhance the quality of the dataset.

To refine the gene expression data further, Soupx (v1.6.2) was used to perform noise correction.^30^ Processed count matrices were loaded into the R environment and analyzed using Seurat (version 5).^31^ To ensure comparability across cells, expression values were subsampled and normalized prior to downstream analysis. Briefly, gene expression data were normalized using SCTransform, followed by identification of highly variable genes. To minimize technical variability arising from independently generated datasets, batch correction and dataset integration were performed using SCT integration, enabling alignment of shared cellular populations prior to downstream analyses. Latent structure in the dataset was captured using principal component–based dimensionality reduction, with the top 30 components retained for subsequent steps. Graph-based community detection was then applied to group cells into transcriptionally distinct populations using a Louvain framework with a resolution parameter of 0.5. Cluster identities were inferred by identifying genes with enriched expression within each group using Seurat’s marker detection workflow, applying a multiple-testing–corrected significance cutoff of 0.05. Low-dimensional embeddings and expression visualizations, including Uniform Manifold Approximation and Projection (UMAP), were generated using Seurat in combination with ggplot2 to illustrate cell population structure and marker gene distributions.

### Cell–Cell Communication Analysis using CellChat

Cell–cell communication was analyzed using the CellChat R package (v2.1.2), which infers intercellular interactions from scRNA-seq data. The analysis was performed by creating a CellChat object that incorporated both the gene expression matrix and annotated cell type metadata. Using curated signaling pathway information, the package identified potential ligand– receptor interactions and highlighted overexpressed genes and signaling events. Communication probabilities were then calculated to determine the intensity and direction of signaling between different cell populations. The resulting interaction networks and prominent signaling pathways were visualized using built-in CellChat plotting tools.

## RESULTS

### Characterization of Cellular Clusters in Osteoarthritis

In the OA group, single-cell sequencing yielded 29,498 cells for further analysis. To investigate the cellular heterogeneity within these cells, we performed a dimensionality reduction analysis. The UMAP plot (**Figure 1A**) illustrates the distinct clusters identified within the sample, highlighting diverse cell populations. We observed 15 distinct clusters, which were annotated as Naïve Tcells, macrophages, B cells, dendritic cells, plasma cells, endothelial cells, chondrocytes, pericytes and neutrophils. Each cluster exhibits a unique spatial organization, indicating varying cellular characteristics and functions. The distinct clusters were assigned to specific cell types based on the expression of recognized marker genes as shown in **Figure 1B**. CellChat analysis revealed global cell–cell communication patterns across all clusters illustrating the relative strength and key intercellular interactions (**Figure 1C**).

**Figure 1.**
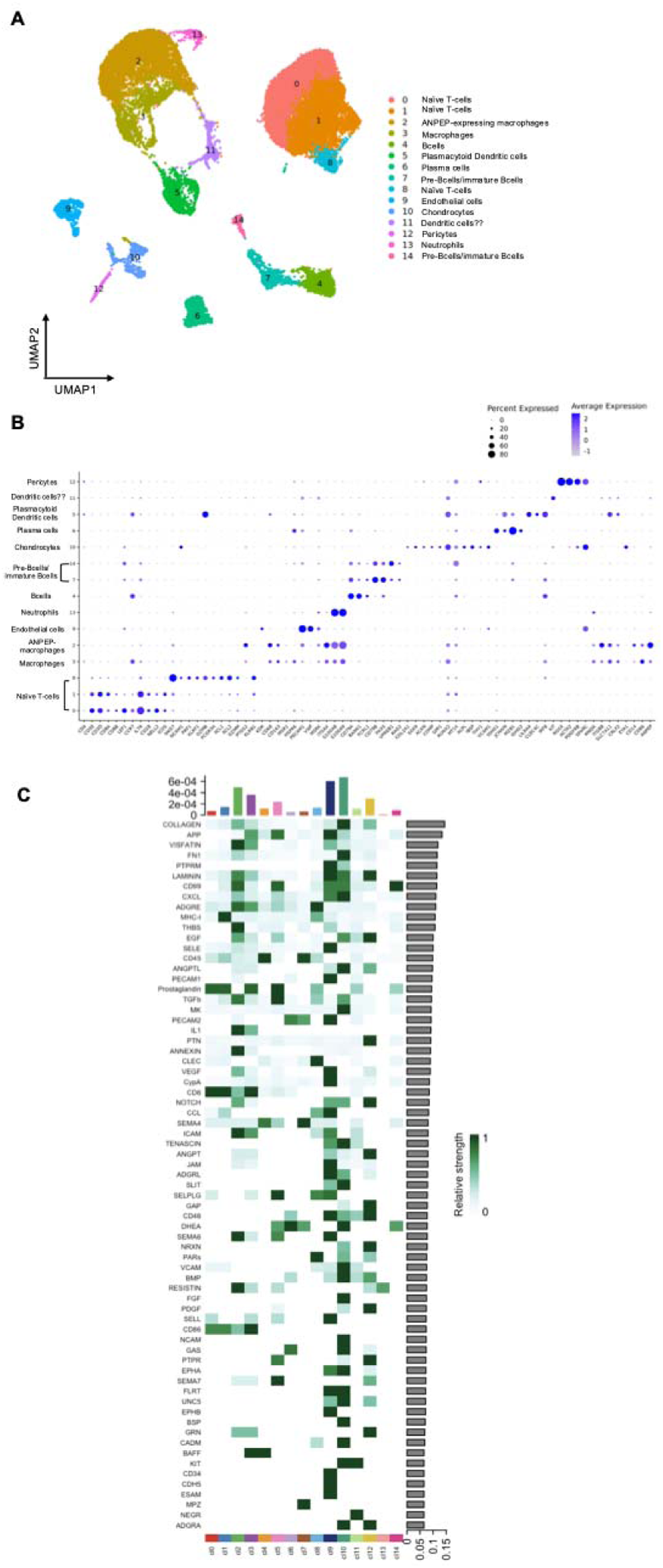
Characterization of cellular heterogeneity in OA. (A) UMAP showing the cell populations identified. (B) Dot plot showing the expression of marker genes in each cluster. Dot sizes are proportional to the percentage of cells in each cluster expressing the indicated gene. (C) Heatmap showing the overall signaling patterns within the different clusters.

### Characterization of Cellular Clusters in Steroid-Induced ONFH

In the steroid-induced ONFH group, single-cell sequencing yielded 8,583 cells for further analysis. The analysis of cellular diversity is illustrated in **Figure 2A**. In total, we identified 20 unique clusters and annotated them as Naïve T cells, endothelial cells, neutrophils, B cells, chondrocytes, macrophages, pericytes, plasma cells and osteoblasts based on marker gene expression (**Figure 2B**). Each cluster displayed a unique arrangement, suggesting distinct functional states and biological roles. Comprehensive analysis using CellChat uncovered the landscape of intercellular communication across all clusters, highlighting dominant signaling routes and quantifying the interaction strength between cell types (**Figure 2C**).

**Figure 2.**
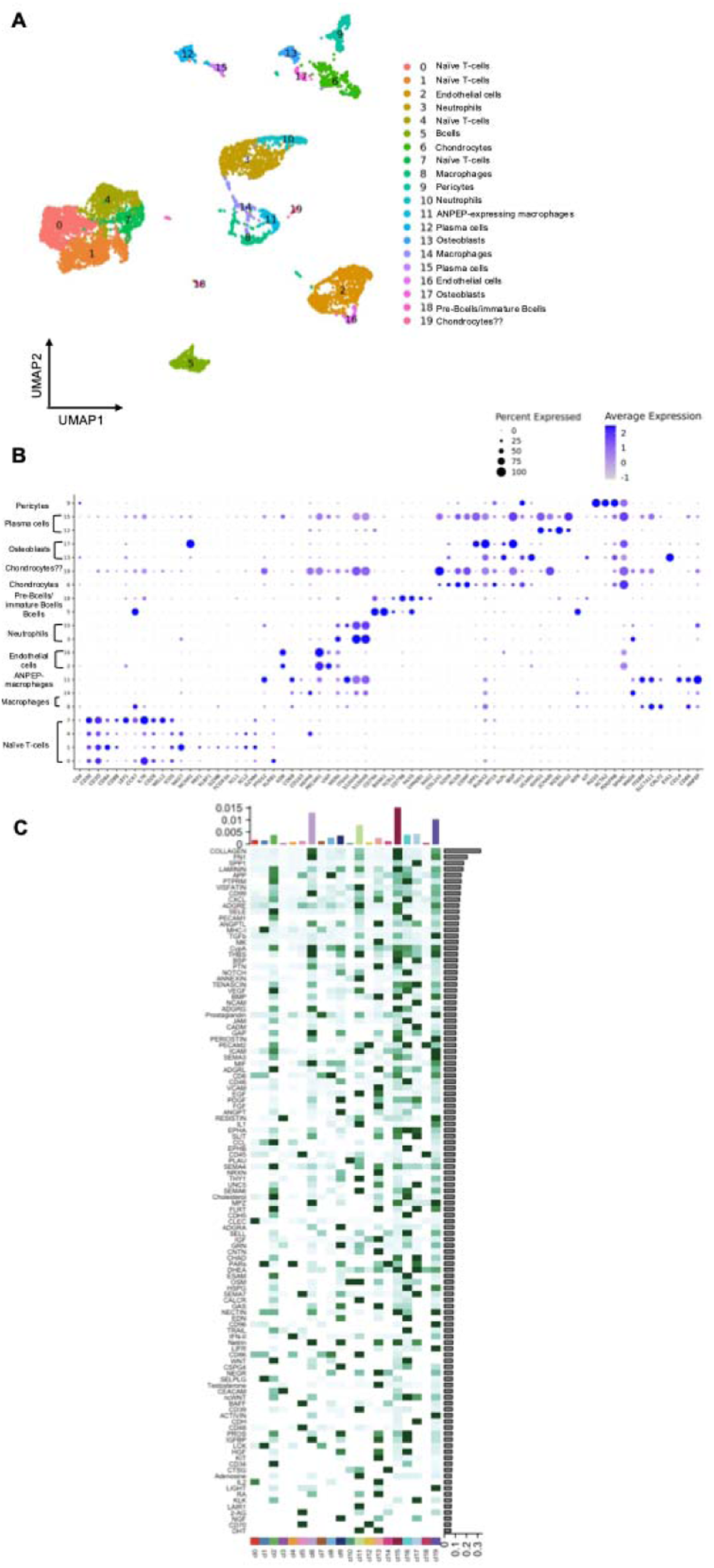
Characterization of cellular heterogeneity in steroid-induced ONFH. (A) UMAP showing the cell populations identified. (B) Dot plot showing the expression of marker genes in each cluster. Dot sizes are proportional to the percentage of cells in each cluster expressing the indicated gene. (C) Heatmap showing the overall signaling patterns within the different clusters.

### Single-Cell Transcriptomics Reveals Distinct Cellular Compositions in Steroid-Induced ONFH and OA

The transcriptomic differences between steroid-induced ONFH and OA were compared by downsampling and combining the datasets containing 17,166 cells (8,583 cells from each sample). Computational analysis of this merged dataset revealed 16 unique clusters with distinct gene expression patterns (**Figure 3**). Major cell populations included T cell subsets, myeloid cells (macrophages, neutrophils, dendritic cells), B cell lineage cells, endothelial cells, and mesenchymal lineage cells (osteoblasts, chondrocytes, fibroblasts, and pericytes) (**Figure 3A-C**).

**Figure 3.**
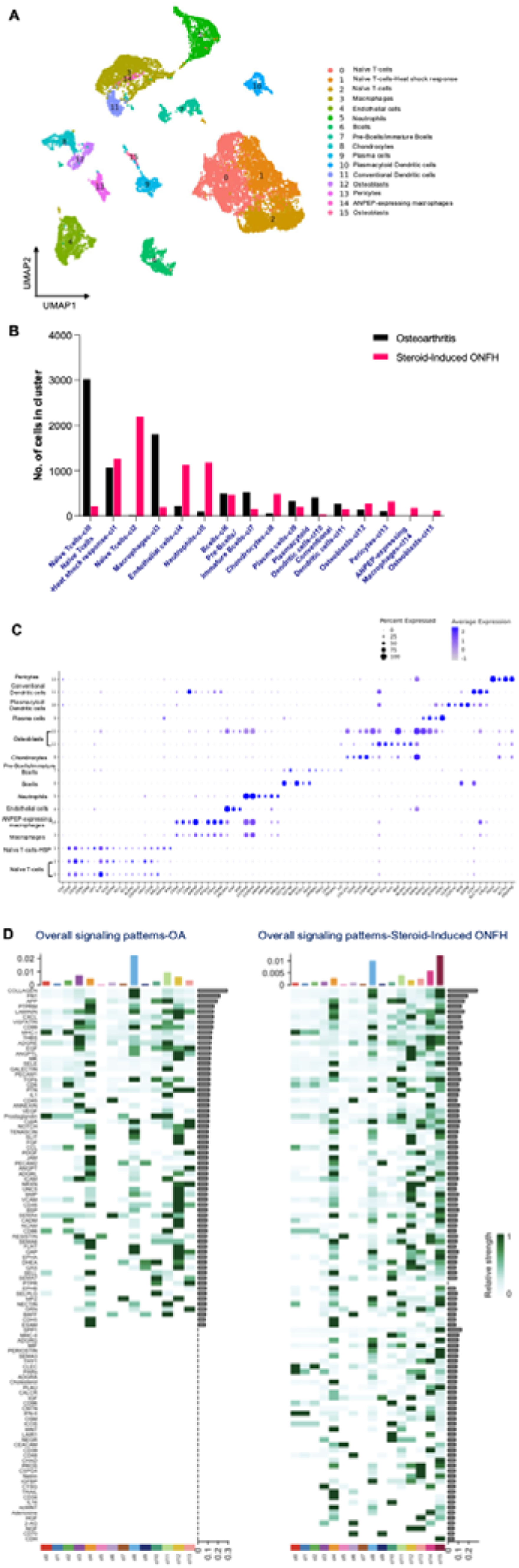
Comparative analysis comparing microenvironment of steroid-induced ONFH to OA (A) UMAP showing the cell populations identified from the combinatorial analysis of 17166 cells (8583 cells from each sample). (B) Bar plot demonstrating the distribution of the population. (C) Dot plot showing the expression of marker genes in each cluster. Dot sizes are proportional to the percentage of cells in each cluster expressing the indicated gene. (D) Heatmap summarizing global signaling patterns across all cell clusters.

Quantitative comparison of cellular compositions revealed notable differences between steroid-induced ONFH and OA. Within T cell subsets, steroid-induced ONFH showed a marked difference in naïve T cell populations compared to OA. A relative reduction in cluster 0 and enrichment of cluster 2, suggesting distinctive functional subsets of naïve T cells in these conditions (**Figure 3B**).

Myeloid cell populations showed condition-specific alterations, with inflammatory macrophages (cluster 3) markedly increased in OA compared to steroid-induced ONFH. In contrast, neutrophils (cluster 5) were enriched in steroid-induced ON compared to OA, pointing to distinct inflammatory mechanisms.

Within stromal cell populations, steroid-induced ONFH showed higher numbers of mesenchymal cells with osteogenic features (clusters 8, 12, and 15) compared to OA. The vascular compartment in ONFH demonstrated increased endothelial cells (cluster 4) and pericytes/smooth muscle cells (cluster 13) compared to OA. Collectively, these features likely reflect greater osteogenic and vascular remodeling in steroid-induced ONFH. Using CellChat, we mapped the intercellular signaling architecture across all clusters, capturing the dynamics and intensity of communication networks among diverse cell populations (**Figure 3D**).

### Single-Cell Transcriptomics Reveals Distinct Cellular Compositions in Steroid-Induced ONFH and Alcohol-Induced ONFH

In the steroid-induced ONFH and alcohol-induced ONFH comparison, the downsampled, combined dataset contained 17,166 cells (8,583 cells from each sample). Unsupervised clustering identified 19 distinct cell clusters (**Figure 4A-C**).

**Figure 4.**
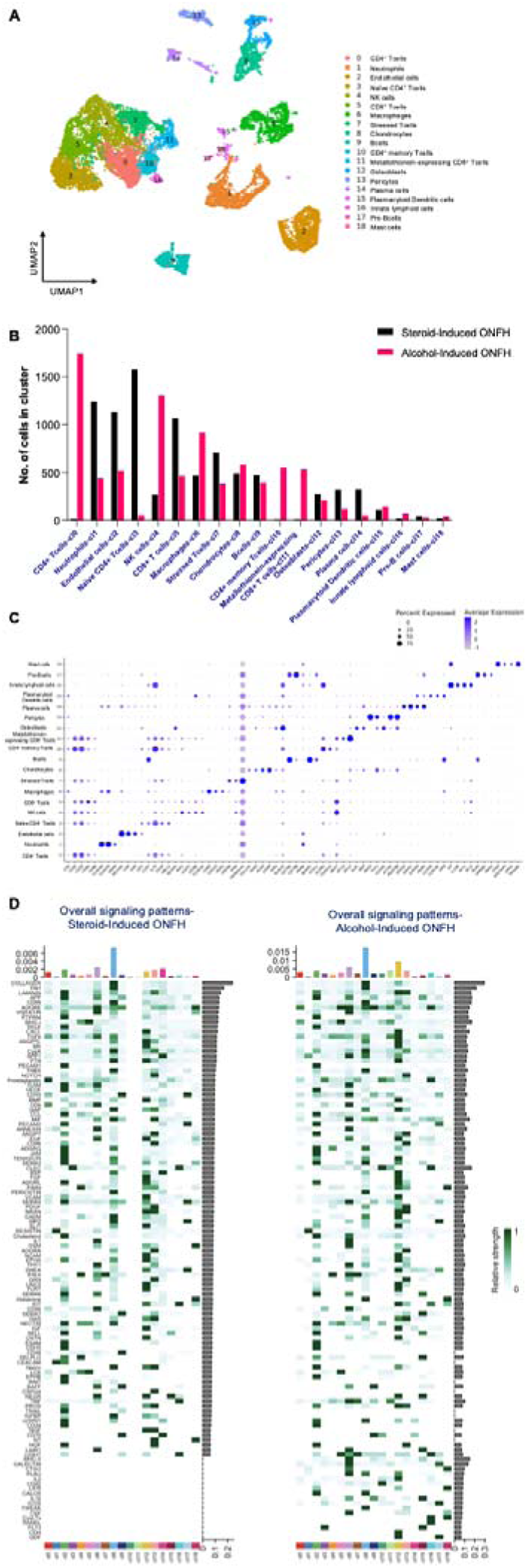
Comparative analysis comparing microenvironment of steroid-induced ONFH to alcohol-induced ONFH (A) UMAP showing the cell populations identified from the combinatorial analysis of 17,166 cells (8583 cells from each sample cells. (B) Bar plot demonstrating the distribution of the population. (C) Dot plot showing the expression of marker genes in each cluster. Dot sizes are proportional to the percentage of cells in each cluster expressing the indicated gene. (D) Heatmap summarizing global signaling patterns across all cell clusters.

When comparing the immune cell composition between steroid-induced and alcohol-induced ONFH, we identified notable differences in innate versus adaptive immune cell proportions. Steroid-induced ONFH showed a higher percentage of innate immune cells (34.2%) compared to alcohol-induced ONFH (23.8% of all cells), primarily due to the marked increase in neutrophils. Conversely, adaptive immune cells represented 18.6% of all cells in steroid-induced ONFH but comprised 29.4% of cells in alcohol-induced ONFH. The resulting innate-to-adaptive immune cell ratio was substantially higher in steroid-induced ONFH (1.84) compared to alcohol-induced ONFH (0.81) or OA (0.62), highlighting fundamental differences in the immune cell landscape between these conditions (**Figure 4B**).

Steroid-induced ONFH showed lower proportions of mature CD4^+^ T cells (cluster 0) and CD4^+^ memory T cells (cluster 10) but higher proportions of naïve CD4^+^ T cells (cluster 3) relative to alcohol-induced ONFH. The overall proportion of CD8^+^ T cells was similar between the two conditions, but the alcohol-induced ONFH group showed a marked enrichment in metallothionein-expressing CD8^+^ T cells (cluster 11), suggesting a distinct cellular stress response mechanism.

Neutrophils (cluster 1) showed markedly higher levels in steroid-induced ONFH compared to alcohol-induced ONFH. This may reflect an ongoing acute inflammatory response in the steroid-induced condition but may also reflect the effects of glucocorticoids on neutrophil lifespan.^32^ Conversely, steroid-induced ONFH samples contained far fewer NK cells (cluster 4) and macrophages (cluster 6), further supporting a distinct steroid-mediated inflammatory response.

Stromal cell populations also showed etiology-specific alterations, with endothelial cells (cluster 2) and pericytes (cluster 13) showing increased numbers in steroid-induced ONFH. We found that the number of chondrocytes and osteoblasts were similar between groups. CellChat analysis revealed that steroid-induced and alcohol-induced ONFH had distinct intercellular communication landscapes, with notable differences in signaling intensity and pathway activity. Several signaling pathways present in one context were entirely absent in the other, indicating condition-specific modes of cell–cell interaction. For example, multiple immunoregulatory pathways were completely absent in steroid-induced ONFH while active in alcohol-induced ONFH, including MHC-II, CD40, CD86, IL2, IL16, LIGHT, and TWEAK signaling. (**Figure 4D**).

### Reclustering T Cell Clusters Reveals Distinct Cellular Subpopulations and Marker Gene Signatures

Given the prevalence of T cell populations across conditions, we performed reclustering of T cell clusters to achieve a higher resolution of the cellular diversity within our downsampled, combined datasets. This analysis allowed us to capture finer subpopulations and functional subsets of T cells that were not fully distinguishable in the initial analysis. Following reclustering, we conducted a comprehensive analysis of marker genes specific to each subpopulation (**Figure 5**).

**Figure 5.**
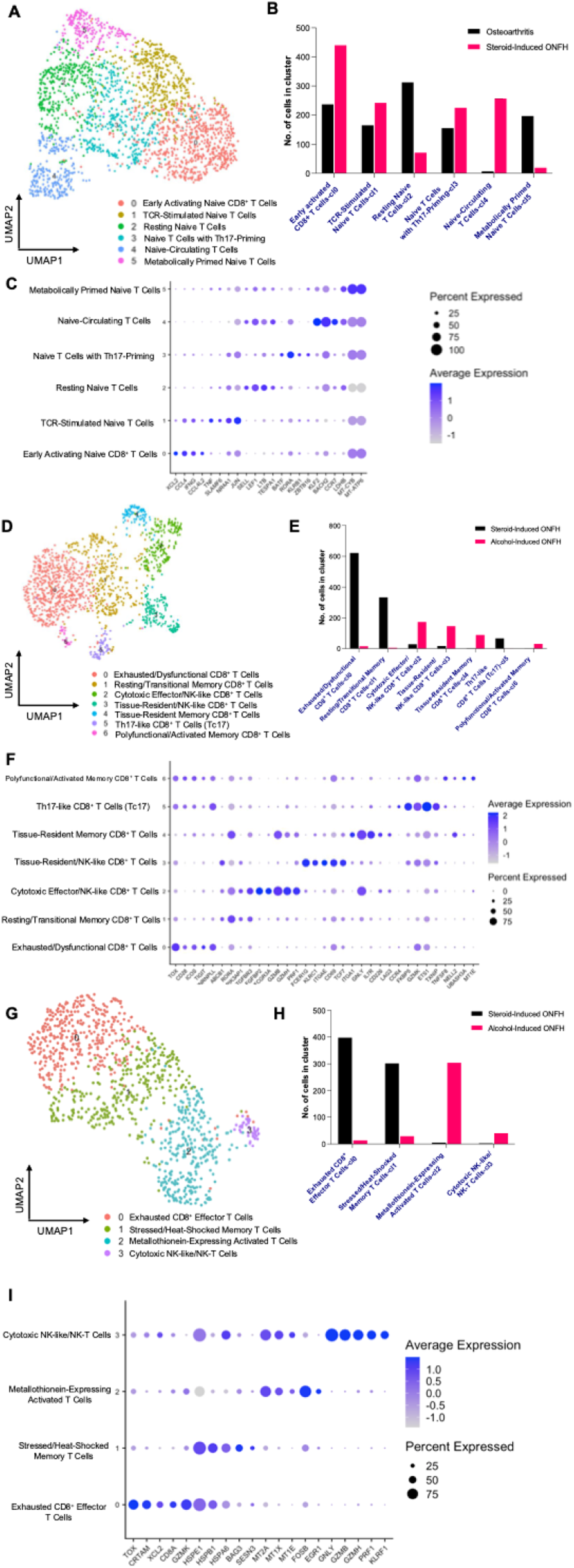
Reclustering and characterizing T cell populations (A) UMAP showing the 6 distinct reclustered T cell populations comparing OA to steroid-induced ONFH. (B) Bar plot demonstrating the distribution of the population comparing OA to steroid-induced ONFH. (C) Dot plot showing the expression of marker genes in each cluster comparing OA to steroid-induced ONFH. Dot sizes are proportional to the percentage of cells in each cluster expressing the indicated gene. D) UMAP showing the 7 distinct reclustered T cell populations comparing steroid-induced to alcohol-induced ONFH. E) Bar plot demonstrating the distribution of the population comparing steroid-induced to alcohol-induced ONFH. F) Dot plot showing the expression of marker genes in each cluster comparing steroid-induced to alcohol-induced ONFH. Dot sizes are proportional to the percentage of cells in each cluster expressing the indicated gene. G) UMAP showing the 4 distinct reclustered T cell populations comparing steroid-induced to alcohol-induced ONFH. H) Bar plot demonstrating the distribution of the population comparing steroid-induced to alcohol-induced ONFH. I) Dot plot showing the expression of marker genes in each cluster comparing steroid-induced to alcohol-induced ONFH. Dot sizes are proportional to the percentage of cells in each cluster expressing the indicated gene.

#### Osteoarthritis and Steroid-induced ONFH T Cell Reclustering Comparisons

In naïve T cell population 1, we identified 10 distinct subclusters with dramatically different distributions between conditions (**Supplemental Figure 1A-C**). OA samples contained abundant effector, memory, cytotoxic, and stress-responsive expressing metallothionein genes (MT1G, MT1X, etc.) and cells with regulatory-like features (cluster 9, FOXP3) suggesting a complex immune landscape involving both pro-inflammatory and regulatory components.

Naïve T cell population 2 (with heat shock response) demonstrated 6 subclusters (**Figure 5A-C**). Early activated CD8^+^ T cells (subcluster 0) and naïve circulating T cells (subcluster 4) were more abundant in ONFH compared to OA. In contrast, resting naïve T cells (subcluster 2) showed over 4-fold higher representation in OA compared to steroid-induced ONFH, while metabolically primed naïve T cells (subcluster 5) were nearly 10-fold more abundant in OA. This pattern suggests steroid-induced ONFH is characterized by dysregulation of naïve T cell homeostasis with a shift toward activation states compared to the T cells residing in a more quiescent state in OA.

We identified 7 subclusters in naïve T cell population 2 (**Supplemental Figure 2A-C**). In contrast to naïve T cell population 1, these T cell subsets were almost exclusively present in steroid-induced ONFH samples but contained similar cell populations including cytotoxic, memory and regulatory T cells. Notably, there was enrichment of CD8^+^primed T cell subpopulations but these cells showed features of exhaustion/stress (e.g. expression of TOX, FKBP5, and MT2A), suggestive of chronic stimulation.

#### Steroid-Induced and Alcohol-Induced ONFH T Cell Reclustering Comparisons

For this comparison, reclustering of CD4^+^ T cells revealed 7 subclusters with minimal contributions from the steroid-induced (**Supplemental Figure 3A-C**). This subcluster displayed substantial heterogeneity, with features of central memory T cells, early activated/stressed T cells, cytotoxic T cells, Th1 cells, Th17 T cells, regulatory T cells, and T cells with innate-like features.

Within the naïve CD4^+^ population, these subclusters were almost entirely expressed by steroid-induced ONFH groups (**Supplemental Figure 4A-C**). The most prevalent features were cell subsets with stress response genes (TXNIP, HSPA1B, HNAJB4) and a Th17-like differentiation signature.

CD8^+^ T cells demonstrated 7 subclusters with marked differences between conditions (**Figure 5D-F**). Exhausted/dysfunctional CD8^+^ T cells (subcluster 0) and resting/transitional memory CD8^+^ T cells (subcluster 1) were dramatically enriched in steroid-induced ONFH, while cytotoxic effector/NK-like CD8^+^ T cells (subcluster 2) and tissue-resident/NK-like CD8^+^ T cells (subcluster 3) were substantially more abundant in alcohol-induced ONFH. This pattern suggests fundamentally different CD8+ T cell responses in these two forms of osteonecrosis, with steroid-induced disease characterized by T cell exhaustion rather than cytotoxic activity.

Stressed T cells were reclustered into 4 distinct subtypes (**Figure 5G-I**). Exhausted CD8^+^ effector T cells (subcluster 0) and stressed/heat-shocked memory T cells (subcluster 1) predominated in steroid-induced ONFH, while metallothionein-expressing activated T cells (subcluster 2) and cytotoxic NK-like or NK-T cells (subcluster 3) were more abundant in alcohol-induced ONFH. This finding highlights the distinct cellular stress response mechanisms in these two etiologies suggesting that steroid-induced ONFH is associated with persistent antigen exposure and T cell dysfunction while alcohol-induced ONFH is associated with an active inflammatory response to oxidative stress.

CD4^+^ memory T cells demonstrated 4 subclusters, with minimal expression in steroid-induced ONFH. Similarly, metallothionein-expressing CD8^+^ T cells were almost exclusively identified in the alcohol-induced ONFH.

Collectively, these reclustering analyses reveal distinctive T cell functional states across conditions, suggesting specific immunological signatures that may contribute to the pathogenesis of ONFH in an etiology-specific manner. The identification of these marker genes and specialized T cell subsets provides a foundation for further functional characterization.

### Summary of Key Cellular Differences that Distinguish Steroid-Induced from Alcohol-Induced ONFH and OA

Our single-cell transcriptomic analysis revealed several distinctive cellular features that differentiate steroid-induced ONFH from both alcohol-induced ONFH and OA (**Table 1**). Among T cell populations, we observed a striking divergence in metallothionein (MT) expression patterns between steroid and alcohol-induced ONFH. While the overall proportion of CD8^+^ T cells was comparable between these conditions, alcohol-induced ONFH demonstrated marked enrichment in metallothionein-expressing CD8^+^ T cells. This distinction is pronounced as MT expression in T cells typically increases in response to inflammatory stimuli, oxidative stress, and cytokines. Though glucocorticoids can directly induce metallothionein expression in T cells, they simultaneously suppress cytokine-mediated MT induction by inhibiting pro-inflammatory cytokine production.^33^ Our data suggests this suppressive effect predominates in steroid-induced ONFH, resulting in reduced MT expression in T cell populations compared to alcohol-induced ONFH.

**Table 1.** Summary of key differences between alcohol-induced and steroid-induced ONFH.

|  | **Myeloid** | **Lymphoid** | **Stromal** |
|---|---|---|---|
| **Steroid-Induced ONFH** | Neutrophil predominant | Increased naïve CD4<sup>+</sup><br>Exhausted/dysfunctional CD8+<br>Increased resting/transitional memory CD8+ T cells<br>Lower CD4+/CD8+ ratio | Increased mesenchymal cells with osteogenic features<br>Increased endothelial cells<br>Similar chondrocyte and osteoblast |
| **Alcohol-Induced ONFH** | Macrophage predominant | Increased metallothionein-expressing T cells<br>Increased cytotoxic NK-like cells | Similar chondrocyte and osteoblast |
| **Osteoarthritis** | Macrophage predominant<br>Decrease neutrophils | Metabolically primed but resting T cells | Decreased endothelial cells |

Further immunophenotypic differences were evident in the CD4^+^/CD8^+^ T cell compartment. Steroid-induced ONFH exhibited fewer mature CD4^+^ T cells and CD4^+^ memory T cells but contained higher proportions of naïve CD4^+^ T cells relative to alcohol-induced ONFH. This dramatic imbalance likely reflects the profound immunosuppressive effects and reduction in CD4^+^ T cell function associated with steroid exposure. The lower CD4/CD8 ratio in steroid-induced ONFH aligns with previous reports in the literature.^34^ Notably, subclustering analysis revealed that naïve CD4^+^ cells in steroid-induced ONFH contained subsets with cytotoxic features, potentially contributing to the inflammatory microenvironment.

The myeloid cell landscape also exhibited condition-specific alterations. OA samples contained increased macrophages compared to steroid-induced ONFH, consistent with the established role of macrophage infiltration as a hallmark of osteoarthritis. In contrast, steroid-induced ONFH demonstrated higher neutrophil numbers compared to both OA and alcohol-induced ONFH. This neutrophilia likely reflects the known effect of glucocorticoids on increasing neutrophil lifespan and may influence neutrophil extracellular trap (NET) formation, which has been associated with microvascular occlusion and local ischemia.^35^ Conversely, alcohol-induced ONFH samples contained greater proportions of NK cells, NKT cells, and macrophages than steroid-induced ONFH, further distinguishing the inflammatory profiles of these etiologies.

The stromal compartment showed etiology-specific alterations as well. Compared to OA, steroid-induced ONFH was associated with a greater number of osteogenic and vascular cells, reflecting a potentially more active bone remodeling microenvironment. Endothelial cells and pericytes were also increased in steroid-induced ONFH compared to alcohol-induced ONFH. Within these vascular populations, we detected expression of inflammatory markers including IL7R, SEMA4D, TNFAIP3, CCR7, and FOXP1, as well as the anti-apoptotic factor BCL2. These findings suggest distinct vascular responses in steroid-induced ONFH, potentially reflecting endothelial activation and altered vascular homeostasis.^8^

Collectively, our findings highlight the complex and etiology-specific cellular landscape in steroid-induced ONFH, characterized by distinctive immune cell compositions, altered T cell maturation states, active but potentially dysfunctional osteogenic cells, and vascular cell expansion with inflammatory markers.

## DISCUSSION

The findings from our single-cell RNA sequencing analysis characterize the cellular and molecular landscape of steroid-induced osteonecrosis of the femoral head (ONFH) and highlight differences relative to osteoarthritis and alcohol-induced ONFH. These observations enhance our understanding of potential etiology-specific cellular and transcriptional alterations and generate hypotheses for future validation studies.

A key discovery in our study is the altered T cell landscape in steroid-induced ONFH compared to OA and alcohol-induced ONFH, revealing previously uncharacterized immune dysregulation. In steroid-induced ONFH, we observed a decrease in the proportion of mature CD4+ T cells but abundant naïve CD4+ T cells. This imbalance likely reflects the impact of steroids on thymic function and T cell development, as glucocorticoids are known to reduce thymic output and impair T cell maturation.^36^ Furthermore, we identified a lack of mature T helper 17 and regulatory T cells in steroid-induced ONFH. When reclustering naïve CD4+ T cells, steroid-induced ONFH was also associated with an elevated ratio of Th17-primed cells compared to Treg-primed cells. A reduction in Treg cells, and an imbalance of Treg/Th17 cells have been previously associated with reduced angiogenesis and increased osteoclastogenesis, suggesting that the altered T cell homeostasis we observed could contribute directly to disease pathogenesis.^34,37–39^ In steroid-induced ONFH there is a shift from quiescent to activated naïve T cell which contrasts with the metabolically primed but resting T cells in OA. These findings suggest that glucocorticoids paradoxically drive T cell activation while simultaneously impairing their function, a phenomenon that may contribute to the failed immune surveillance and impaired tissue repair characteristic of ONFH.

In addition to differences in T cell maturation, we also observed differences in T cell functional states. Steroid-induced ONFH was characterized by dysfunctional/exhausted CD8+ T cells, and CD8+ T cells with reduced metallothionein (MT) expression compared to alcohol-induced ONFH, reflecting distinct immune responses between these conditions. This finding is significant as MTs serve as important antioxidant and anti-inflammatory mediators that protect against oxidative stress.^40–42^ The suppression of MT expression in steroid-induced ONFH suggests increased vulnerability to oxidative damage, which aligns with previous studies demonstrating that glucocorticoid-induced oxidative stress contributes to osteoblast dysfunction.^43,44^ In contrast, alcohol-induced ONFH showed robust MT expression, potentially representing a compensatory protective mechanism against alcohol-mediated oxidative injury, consistent with reports that MTs can attenuate hepatic and neural oxidative damage from ethanol toxicity.^45,46^ This deficit in protective mechanisms could explain the more rapid progression of disease often observed in steroid-induced versus alcohol-induced ONFH, as noted in prior histopathological studies.^47,48^

The striking neutrophil population in steroid-induced ONFH compared to both OA and alcohol-induced ONFH represents another novel finding with pathogenic implications. While seemingly counterintuitive given the anti-inflammatory properties of glucocorticoids, this observation is consistent with clinical studies reporting neutrophilia in patients receiving glucocorticoid therapy, though the vascular dysfunction characteristic of ONFH may lead to neutrophils in the perivascular space.^49^ Mechanistically, steroids prolong neutrophil survival by inhibiting apoptosis while simultaneously dampening their functional capacity.^50^ The accumulation of functionally impaired neutrophils may contribute to pathogenesis through formation of neutrophil extracellular traps (NETs), which have been implicated in vascular occlusion and thrombosis.^35^ Nonokawa et al. demonstrated increased NETs in femoral head samples from steroid-induced ONFH patients, suggesting that steroid-mediated neutrophil accumulation could directly contribute to microvascular occlusion and subsequent ischemic injury in the femoral _head._51,52

The vascular compartment alterations we identified in steroid-induced ONFH, specifically the expansion of endothelial cells and pericytes compared to alcohol-induced ONFH, suggest a distinct vascular response. This finding contrasts with the conventional understanding that reduced vascularity drives ONFH pathogenesis.^5^ However, recent evidence indicates that glucocorticoids can induce endothelial dysfunction and activation rather than simply reducing vessel density.^53,54^ The enrichment of inflammatory markers (i.e. IL7R, SEMA4D, TNFAIP3) in these expanded vascular populations suggests a state of endothelial activation, which could impair vascular function despite increased cell numbers. This aligns with findings from Vogt et al. who demonstrated that glucocorticoids can induce endothelial cell activation and glycocalyx degradation, leading to increased permeability and impaired microcirculation.^55^

In contrast to steroid-induced ONFH, alcohol-induced ONFH demonstrated a distinctive cellular signature characterized by robust metallothionein expression in T cells, increased NK and NKT cells, and macrophage predominance. This immune profile is consistent with the known effects of chronic alcohol exposure on promoting pro-inflammatory immune responses, particularly through activation of innate immunity.^56^ The contrasting cellular landscapes between steroid and alcohol-induced ONFH highlight divergent pathogenic mechanisms despite similar clinical presentations, suggesting that potential therapeutic approaches may need to be tailored to the specific etiology.

The translational implications of our findings are significant. The identification of exhausted/dysfunctional CD8^+^ T cells and impaired CD4^+^ T cell maturation in steroid-induced ONFH suggests potential therapeutic opportunities through immune modulation. Recent studies have shown that targeting T cell exhaustion can improve outcomes in various inflammatory conditions, and our preliminary findings suggest similar approaches might benefit patients with steroid-induced ONFH.^46^ Additionally, the expansion of vascular cells with inflammatory phenotypes presents another potential therapeutic target, as modulating endothelial activation could potentially preserve microvascular function and mitigate ischemic injury.

There are important limitations of our study. The relatively small sample size for each condition may have failed to capture the spectrum of cellular variations within each disease state. In addition, given the limited number of biological donors included in this study, donor-specific variability cannot be fully excluded and donor-level statistical interpretation remains limited. Therefore, the findings should be considered exploratory and hypothesis-generating, warranting validation in larger independent cohorts. Second, we were unable to compare our experimental samples to true disease-free controls, as all femoral head specimens were obtained from patients undergoing total hip arthroplasty. While using OA as a comparator allows us to distinguish ONFH-specific changes from general end-stage hip pathology, both conditions share degenerative features that may mask some disease-specific alterations. Third, our cross-sectional design captures only a single timepoint in the disease process -specifically, end-stage disease requiring surgical intervention. This prevents us from understanding the temporal dynamics of cellular changes and identifying which alterations are causative versus compensatory. We had originally planned to include patients with early-stage ONFH undergoing femoral head-preserving procedures such as core decompression; however, the limited quantity and quality of tissue obtained from these minimally invasive procedures precluded single-cell analysis. Fourth, while tissue was collected from multiple zones across the coronal cross-section of the femoral head, including the necrotic zone, sclerotic zone, healthy margin, and subchondral bone, we were unable to perform zone-specific analyses; future spatially resolved transcriptomic approaches would allow cellular characterization at higher anatomical resolution. Fifth, although batch-correction strategies were applied to reduce technical variation between externally sourced datasets, residual batch effects associated with differences in sample processing and sequencing conditions cannot be completely excluded. Sixth, the available demographic information and past medical history for all samples are provided in Supplementary Table 1; however, the limited cohort size precluded systematic evaluation of the effects of clinical heterogeneity, medication exposure, and disease stage on the observed findings. Finally, while we identified distinct cellular populations and gene expression patterns, functional validation through *in vitro* and *in vivo* experiments remains necessary to confirm the mechanistic roles of these findings. Future studies should address these limitations through larger cohorts with longitudinal sampling, inclusion of early-stage disease specimens, and integration of multi-omics approaches to provide a more comprehensive understanding of ONFH.

## CONCLUSION

In summary, our single-cell transcriptomic analysis has uncovered distinct cellular and molecular features that differentiate steroid-induced ONFH from OA and alcohol-induced ONFH. These findings highlight the complex interplay between immune dysregulation, vascular activation, and altered bone homeostasis. By elucidating etiology-specific cellular and molecular changes, our study expands current understanding of steroid-induced ONFH and provides a resource for future studies aimed at validating potential therapeutic targets.

## FUNDING

This study was partially supported and funded by a research grant from the Orthopaedic Research and Education Foundation (OREF). This study was also partially supported by a gift from the Chau Family Charitable Foundation.

The authors would like to thank the USC Flow Cytometry Facility and the USC Molecular Genomics Core for their assistance with this project.

## AUTHOR CONTRIBUTIONS

Conceptualization, AS, MCG, JAB, CKM, DE, JRL; methodology, AS, MCG, JAB, CKM, JRB, OS, DE, JRL; software, AS; validation. AS; formal analysis, AS; investigation, AS, MCG, JAB, CKM, JRB, FH, ELZ, TS, OS ; resources, JRL; data curation, AS, MCG, JAB, CKM, JRB, FH,,

ELZ, TS, OS; writing—original draft preparation, AS, MCG, JAB; writing—review and editing, AS, MCG, JAB, CKM, JRB, FH, ELZ, TS, DE, JRL; visualization, AS.; supervision, AS, DE, JRL; project administration, JRL.; funding acquisition, JRL. All authors have read and reviewed the manuscript.

## DATA AVAILABILITY STATEMENT

The single cell RNA sequencing datasets generated during the current study are available in GEO database under the accession number: GSE316957

