## Supplementary File for "Single-Cell RNA Sequencing Reveals Distinct Cellular and Molecular Features of Steroid-Induced Osteonecrosis of the Femoral Head Compared to Alcohol-Induced Osteonecrosis and Hip Osteoarthritis"

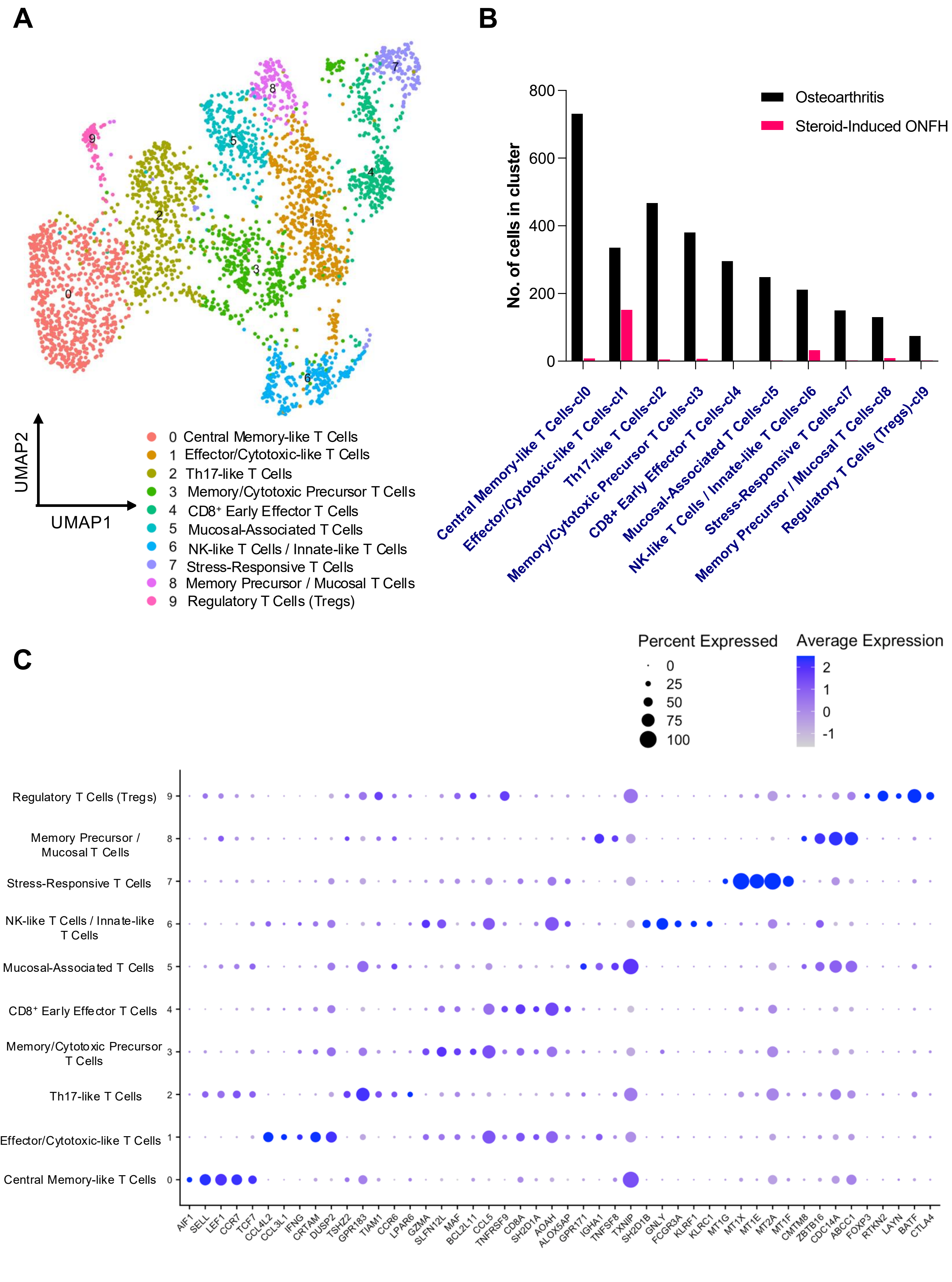


**Supplementary Figure 1.** (A) UMAP showing the cell populations identified from naïve T cell population 1 in osteoarthritis and steroid-induced ONFH comparisons (B) Bar plot demonstrating the distribution of the population. (C) Dot plot showing the expression of marker genes in each cluster. Dot sizes are proportional to the percentage of cells in each cluster expressing the indicated gene.

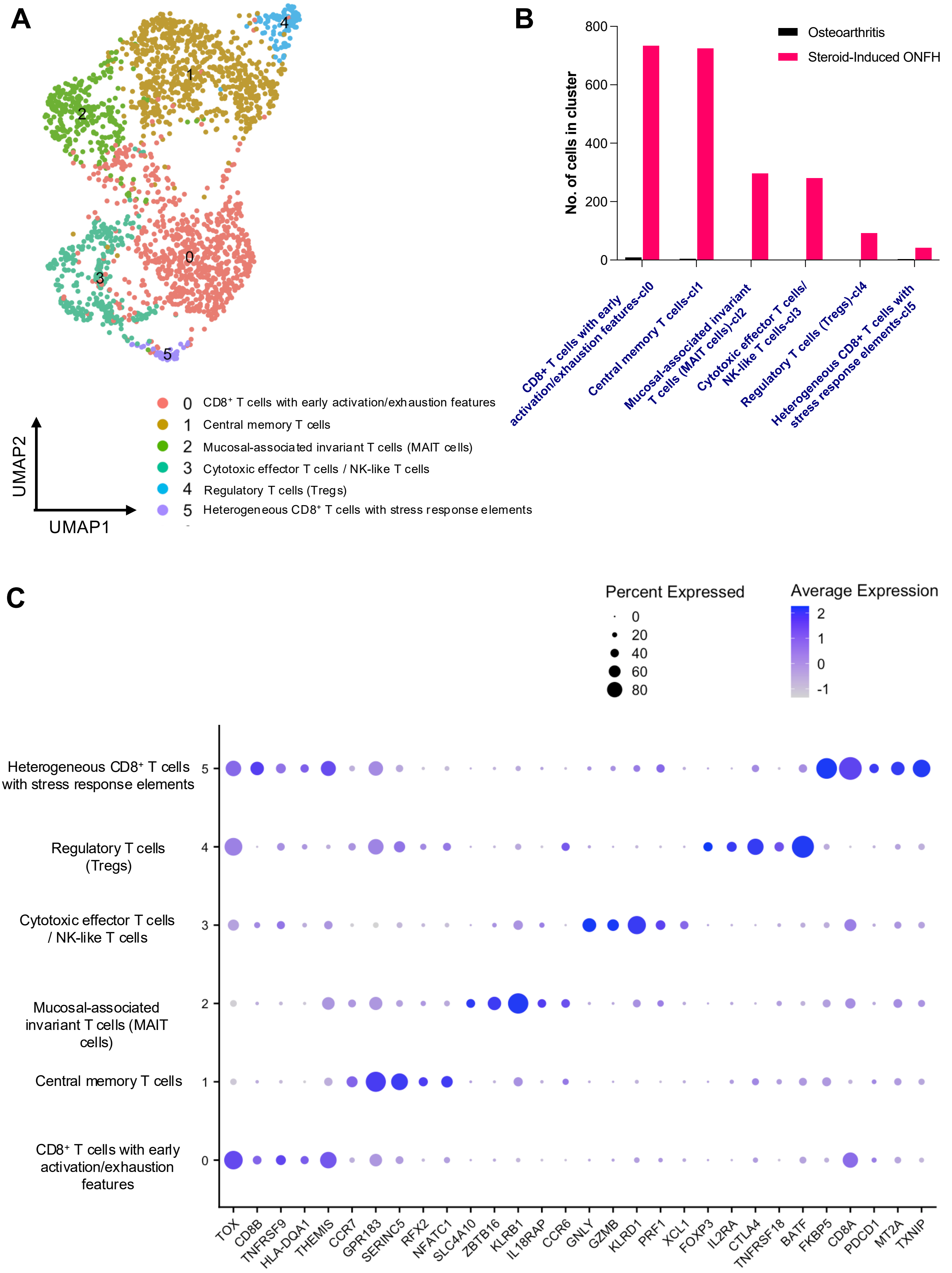


**Supplementary Figure 2.** (A) UMAP showing the cell populations identified from naïve T cell population 2 in osteoarthritis and steroid-induced ONFH comparisons (B) Bar plot demonstrating the distribution of the population. (C) Dot plot showing the expression of marker genes in each cluster. Dot sizes are proportional to the percentage of cells in each cluster expressing the indicated gene.

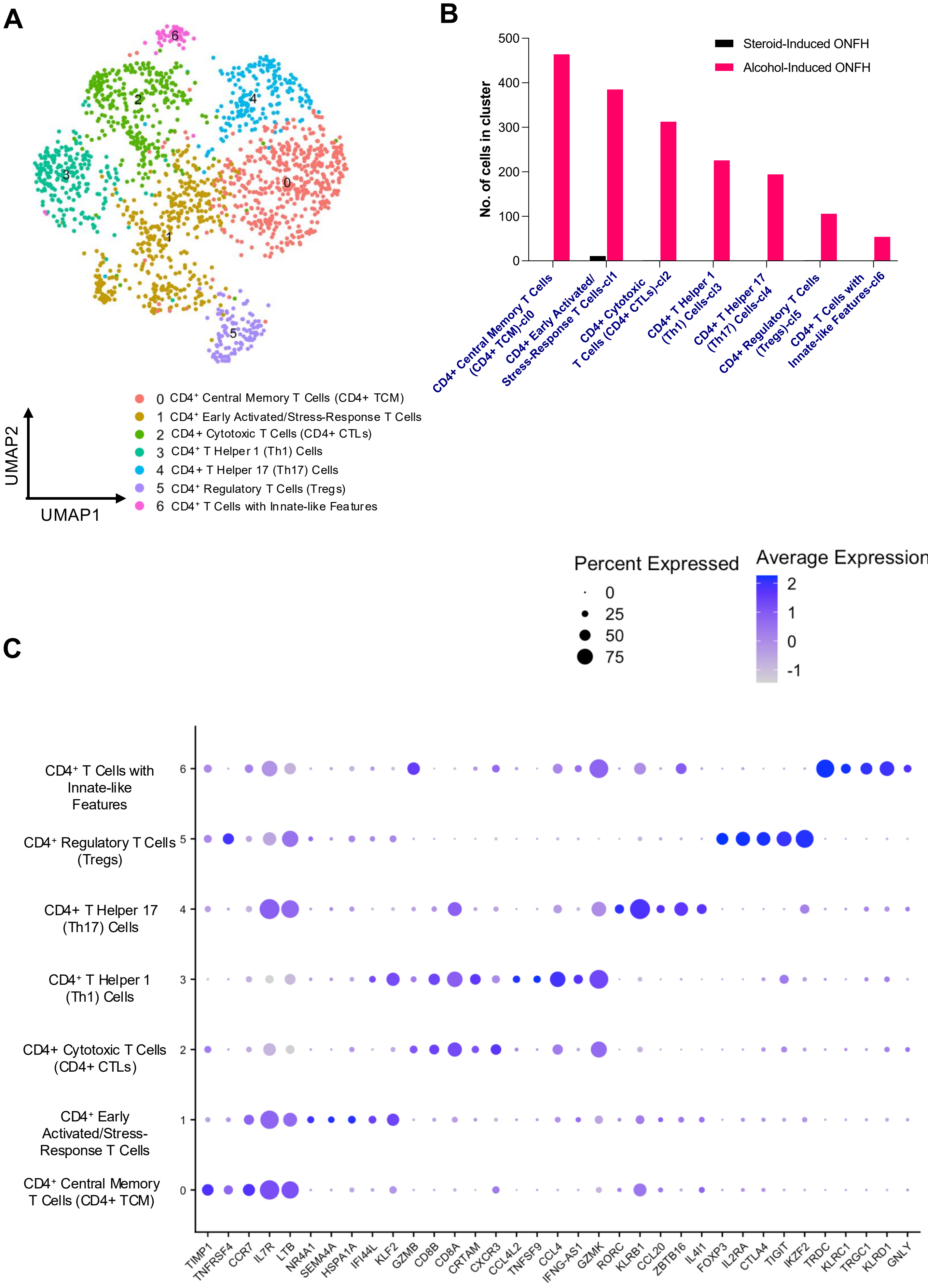


**Supplementary Figure 3.** (A) UMAP showing the cell populations identified from CD4+ T cells in steroid-induced ONFH and alcohol-induced ONFH comparisons (B) Bar plot demonstrating the distribution of the population. (C) Dot plot showing the expression of marker genes in each cluster. Dot sizes are proportional to the percentage of cells in each cluster expressing the indicated gene.

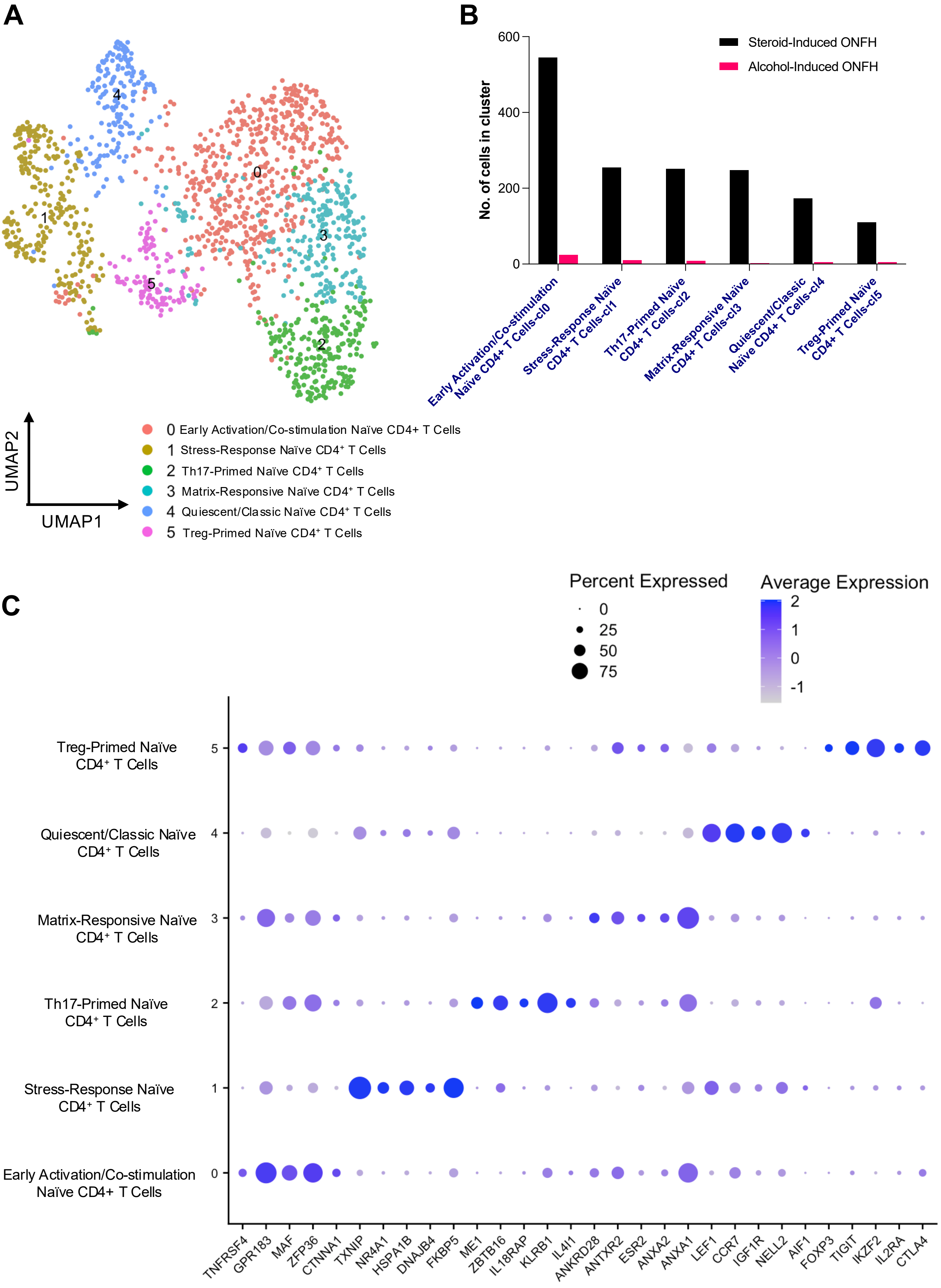


**Supplementary Figure 4.** (A) UMAP showing the cell populations identified from naïve CD4+ population in steroid-induced ONFH and alcohol-induced ONFH comparisons (B) Bar plot demonstrating the distribution of the population. (C) Dot plot showing the expression of marker genes in each cluster. Dot sizes are proportional to the percentage of cells in each cluster expressing the indicated gene.

| Group | Age | Gender | Race | Procedure | Medical History |
|---|---|---|---|---|---|
| Steroid-Induced ONFH | 37 | M | Asian | R THA | HTN, COVID-19 Pneumonia requiring corticosterioids, bilateral ONFH |
| Steroid-Induced ONFH | 37 | M | Asian | L THA | HTN, COVID-19 Pneumonia requiring corticosterioids, bilateral ONFH |
| Osteoarthritis | 49 | M | Caucasian | R THA | HTN, Irregular Heart Beat |
| Osteoarthritis | 49 | F | Black | L THA | HTN, Sickle Cell Trait, Sjogren |
| Osteoarthritis | 60 | F | Asian | L THA | HTN |

S**upplementary Table 1.** Demographic and past medical history of steroid-induced ONFH and OA samples.
